# PHYTOCHROME INTERACTING FACTORs Trigger Environmentally Responsive Chromatin Dynamics

**DOI:** 10.1101/826842

**Authors:** Björn C. Willige, Mark Zander, Amy Phan, Renee M. Garza, Shelly A. Trigg, Yupeng He, Joseph R. Nery, Huaming Chen, Joseph R. Ecker, Joanne Chory

**Author notes:** These authors contributed equally (Björn C. Willige, Mark Zander). Author for correspondence: Joseph R. Ecker and Joanne Chory.

## Abstract

The pivotal interplay between light receptors and PHYTOCHROME INTERACTING FACTORs (PIFs) serves as an essential regulatory hub that perceives and integrates environmental cues into the plant’s transcriptional networks. A critical control component of environmentally-responsive gene networks is the histone variant H2A.Z which provides transcriptional plasticity and prevents undesired gene activation. However, the functional relationship between PIF transcription factors and H2A.Z is only poorly understood. Here, we describe a genomic approach that utilizes the rapid and reversible light-mediated manipulation of PIF7 activity to visualize PIF7 DNA binding and H2A.Z occupancy kinetics. Strikingly, PIFs shape the H2A.Z landscape in a light quality-dependent manner. In addition, we observed that PIFs initiate H2A.Z eviction through direct interaction with EIN6 ENHANCER (EEN), a subunit of INO80 chromatin remodeling complex. These studies uncover a previously unknown PIF-INO80 regulatory module controlling plant growth in response to rapid environmental changes.

**One-sentence summary:** A PIF-INO80 module controls light quality-dependent H2A.Z dynamics.

## Introduction

The environmental cues light and temperature control all aspects of plant growth and are perceived by the phytochrome (phy) family of photoreceptors (*1, 2*). Phytochromes regulate growth by directly modulating the activity of the PIF transcription factors (TFs), which govern a massive environmentally-responsive transcriptional network (*3*). The evolutionary conserved histone variant H2A.Z has emerged as a key regulator of plant environment interactions (*4–7*) and has been postulated to function as temperature sensor in *Arabidopsis* (*5*). As a result of its eviction from gene bodies, H2A.Z chromatin dynamics enable rapid transcriptional responses to various environmental stimuli in *Arabidopsis* (*6–8*). While the current model suggests that H2A.Z occupancy modulates PIF4 binding, this supposition is solely based on the ambient temperature-induced binding of PIF4 to the *FLOWERING LOCUS T* (*FT*) gene (*4*).

## Results

Here, we utilized the light quality-dependent manipulation of PIF7 to visualize H2A.Z dynamics on a genome-wide scale. Simulated competing vegetation, a light condition where a reduced red (R) to far-red (FR) ratio (low R:FR) induces the shade avoidance response and therefore growth (*9*), was used as our environmental cue. We utilized this assay to assess the effects of low R:FR-induced hypocotyl elongation in *pif* multiple mutant combinations (up to a *pif1 pif3 pif4 pif5 pif7* quintuple mutant (*pif13457*), as well as in *phyB* mutants (Fig. 1A, B). Our analysis confirmed that *PIF7* was the major growth regulator followed by *PIF4* and *5*, while mutations in *PIF1* and *3* had only marginal influence on hypocotyl elongation in low R:FR light (Fig. 1A, B). Overexpressed PIF7 is rapidly dephosphorylated after low R:FR light exposure and stimulates hypocotyl growth already in white light (WL) (*10*). To create a more switch-like response of low R:FR-induced PIF7 activity, we introduced a MYC-tagged PIF7 genomic fragment into the *pif457* triple mutant (*pif457 PIF7:PIF7:4xMYC*) that fully complemented the hypocotyl elongation phenotypes of *pif457* and *phyB pif457* mutants (Fig. S1A, B). Immunoblot analyses revealed that PIF7 is phosphorylated throughout a long day (LD) cycle in WL-grown seedlings and was most abundant at Zeitgeber time (ZT) 4. However, during the night PIF7 levels were relatively low and less phosphorylated (Fig. 1C, Fig. S1C). Upon exposure to low R:FR light, dephosphorylation of PIF7 was observed at all time points, and also in WL-exposed *phyb pif457 PIF7:PIF7:4xMYC* mutant lines (Fig. 1C, D, Fig. S1C). These results demonstrate the phyB-dependent control of PIF7 phosphorylation and activity (Fig. 1C, D, Fig. S1B).

**Fig. 1.**
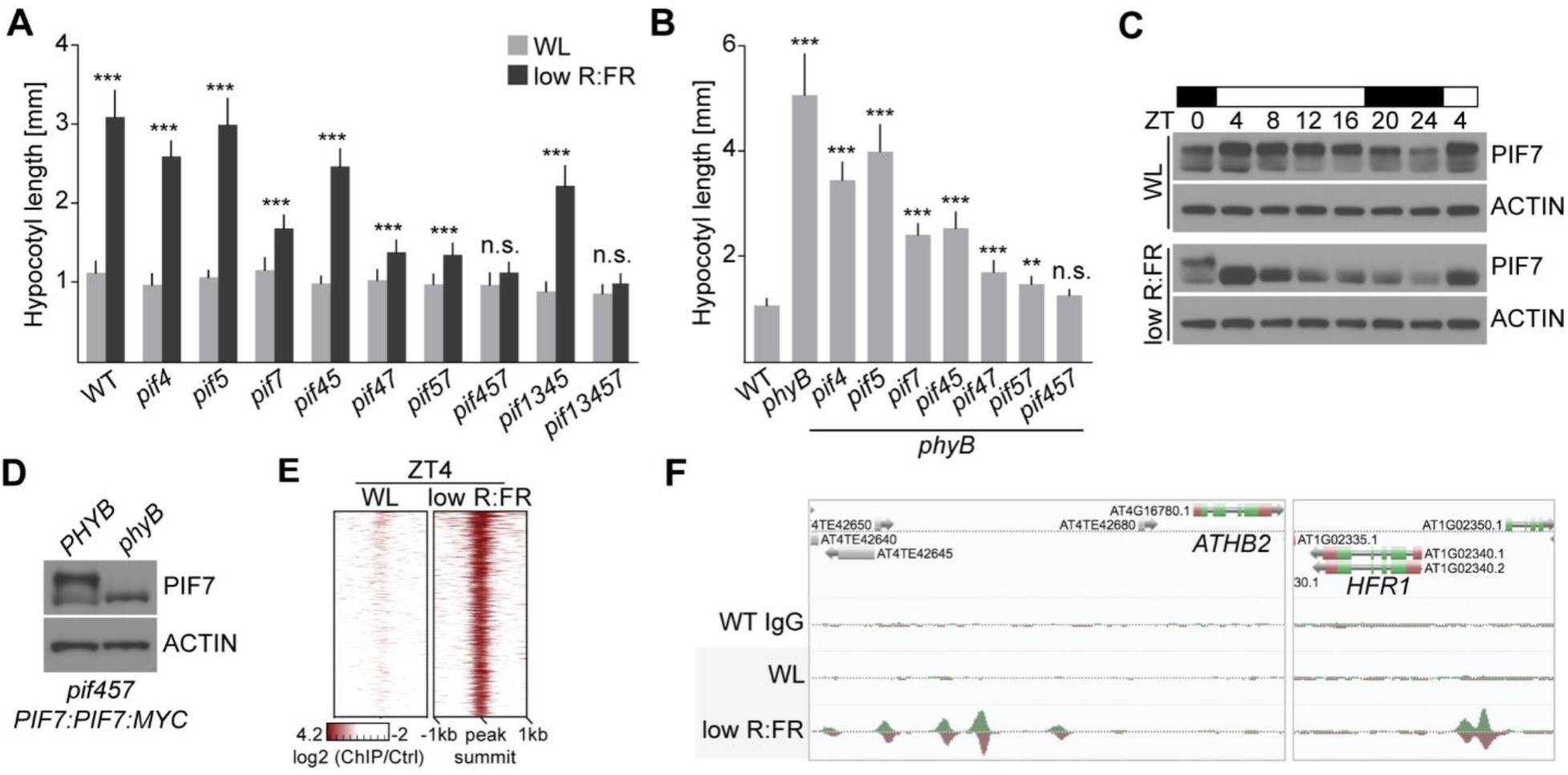
Low R:FR exposure induces dephosphorylation and subsequent genome-wide binding of PIF7. (A) Hypocotyl length measurements of WT and *pif* mutant seedlings grown in WL or in responses to low R:FR. Error bars for the hypocotyl length represent SD (n ≥ 16). Stars denote statistically significant differences between WL and low R:FR for the respective genotypes (Two-way ANOVA, Tukey’s multiple comparisons test, n.s. P > 0.05 * P ≤ 0.05, ** P ≤ 0.01, *** P ≤ 0.001). (B) Hypocotyl length measurements of WT, *phyB* and *phyB pif* mutant combinations grown in WL. Error bars for the hypocotyl length represent SD (n ≥ 20). Stars denote statistically significant differences between WT and mutants (One-way ANOVA, Tukey’s multiple comparisons test, n.s. P > 0.05 * P ≤ 0.05, ** P ≤ 0.01, *** P ≤ 0.001). (C) Immunoblot analysis of *pif457 PIF7:PIF7:4xMYC* in LD. 6-day-old seedlings continued to grow in WL or where exposed to low R:FR at ZT0. (D) Immunoblot analysis of 6-day-old WL-grown *pif457PIF7:PIF7:4xMYC* and *phyB pif457 PIF7:PIF7:4xMYC* seedlings. (E) Heatmap visualizes the low R:FR-induced DNA binding (log_2_ fold change) of PIF7 in *pif457 PIF7:PIF7:4xMYC* seedlings at ZT4. PIF7 DNA binding is shown for the top 500 binding sites that display the strongest low R:FR-induced PIF7 binding at ZT4. PIF7 occupancy is shown from 1 kb upstream to 1 kb downstream of the 500 strongest PIF7 binding events. (F) AnnoJ genome browser screenshot visualizes PIF7 binding at the *ATHB2* and *HFR1* gene. All tracks were normalized to the respective sequencing depth.

Next, we investigated the impact of low R:FR exposure on the genome-wide DNA binding of PIF7 at ZT4 and revealed 1052 significant binding events using ChIP-seq (Table S1). As previously reported for other PIFs (*11–14*), PIF7 preferentially bound to G-boxes (CACGTG) of genes that are predominantly involved in auxin signaling (Fig. S1D, F). Interestingly, less than 20% of PIF7 binding events occurred in proximal (1000 bp) promoter regions (Fig. S1E). In contrast, only very weak PIF7 DNA binding could be detected in WL conditions (Fig. 1E, F, Fig. S1G), supporting the idea that PIF7 dephosphorylation mediates DNA binding. Among the low-R:FR light-dependent genes targeted by PIF7 were the well-studied hypocotyl growth regulators *ATHB2* (*ARABIDOPSIS THALIANA HOMEOBOX PROTEIN 2*) and *HFR1* (*LONG HYPOCOTYL IN FAR-RED*) (*15, 16*) (Fig. 1F).

Next, we examined the impact of low R:FR exposure on the H2A.Z landscape in WT seedlings in a LD time course experiment (ZT0 - ZT20). In line with previous studies (*8, 17, 18*), H2A.Z was specifically enriched in gene bodies of up to 20411 protein-coding genes (Fig. S2A, Table S2). The H2A.Z landscape was highly responsive to low R:FR treatment with up to 1196 genes at dusk (ZT16) showing a low R:FR-induced H2A.Z eviction (Fig. S2B, C, Table S3). Only minimal overlap (ZT0 vs ZT8: 30 genes; ZT0 vs ZT16: 31 genes; ZT8 vs ZT16: 60 genes) was found between genes that display a low R:FR-induced H2A.Z eviction at ZT0, ZT8 and ZT16 (Fig. S2B, C) indicating that the diurnal cycle and low R:FR exposure affect global H2A.Z occupancy as demonstrated by the H2A.Z cycling pattern observed for the flowering regulator *CONSTANS-like 5* (*COL5*) (*19*) (Fig. S2D, E).

To further explore the interplay between PIFs and H2A.Z occupancy, WT seedlings grown in constant light, were exposed to low R:FR light for up to 2 hours, which was then followed by an additional WL phase to capture H2A.Z recovery dynamics (Fig. S3A). Transcriptomes were profiled using RNA-seq and global H2A.Z occupancy was monitored via ChIP-seq. We found that low R:FR-activated gene expression was accompanied by H2A.Z eviction, whereas R:FR-induced gene repression did not coincide with H2A.Z incorporation (Fig. S3B, Table S4). This low R:FR-induced H2A.Z eviction pattern is reversible and disappeared after 2 hours of WL recovery (Fig. S3C). Interestingly, the gene that displayed the most dynamic H2A.Z pattern was *ATHB2* (Fig. S3D, Table S5), a homo-domain transcription factor that mediates growth responses in shade (*15*). After only 30 min of low R:FR exposure H2A.Z levels at *ATHB2* were sharply depleted but rapidly recovered to normal levels during the WL recovery phase (Fig. S3D). A similar H2A.Z pattern was observed for a set of genes (top50, Table S5) with the strongest low R:FR-induced H2A.Z eviction (Fig. S3D). To test the role of PIFs in facilitating low R:FR-induced H2A.Z eviction, we profiled H2A.Z occupancy over time (15 min, 30 min, 2 hours) in *pif457* triple mutants. Strikingly, low R:FR-induced H2A.Z eviction was strongly compromised in low R:FR light-exposed *pif457* seedlings (Fig. 2A, Fig. S4A, Table S6) suggesting that PIFs regulate low R:FR-induced H2A.Z eviction. Specifically, exposure to low R:FR light for only 15 min was sufficient to trigger PIF-dependent H2A.Z eviction (Fig. 2A, Fig. S4A), suggesting that PIF7 DNA/G-box binding might occur even earlier. To test this possibility, PIF7 DNA binding was examined at even shorter low R:FR light exposure times (5, 10 and 30 min) and H2A.Z occupancy and mRNA expression were monitored in parallel. Immunoblot analyses revealed that only 5 min exposure to low R:FR light was sufficient to trigger PIF7 de-phosphorylation (Fig. 2B) which was correlated with a rapid increase in PIF7 DNA binding (Fig. 2C-D, Fig. S4B, C, Table S7). We next assessed H2A.Z occupancy on a set of 20 genes (PIF7 core genes) that show strongest PIF7 binding and activated expression (Table S8). This analysis revealed similar temporal kinetics of low R:FR-induced PIF7 binding and gene body H2A.Z eviction at the PIF7 core gene set which also includes *ATHB2* (Fig. 2D-F). The observation that H2A.Z eviction precedes changes in gene expression indicated that chromatin remodeling is not a consequence of transcriptional activation (Fig. 2E, F). Although our experimental resolution was not able to unravel temporal DNA binding of PIF7 from H2A.Z eviction, our observation of a compromised low R:FR-induced H2A.Z eviction in *pif457* triple mutants (Fig. 2A, Fig. S4A) supports a scenario where low R:FR-induced PIF7 binding is a prerequisite for H2A.Z eviction and subsequent activation of gene expression.

**Fig. 2.**
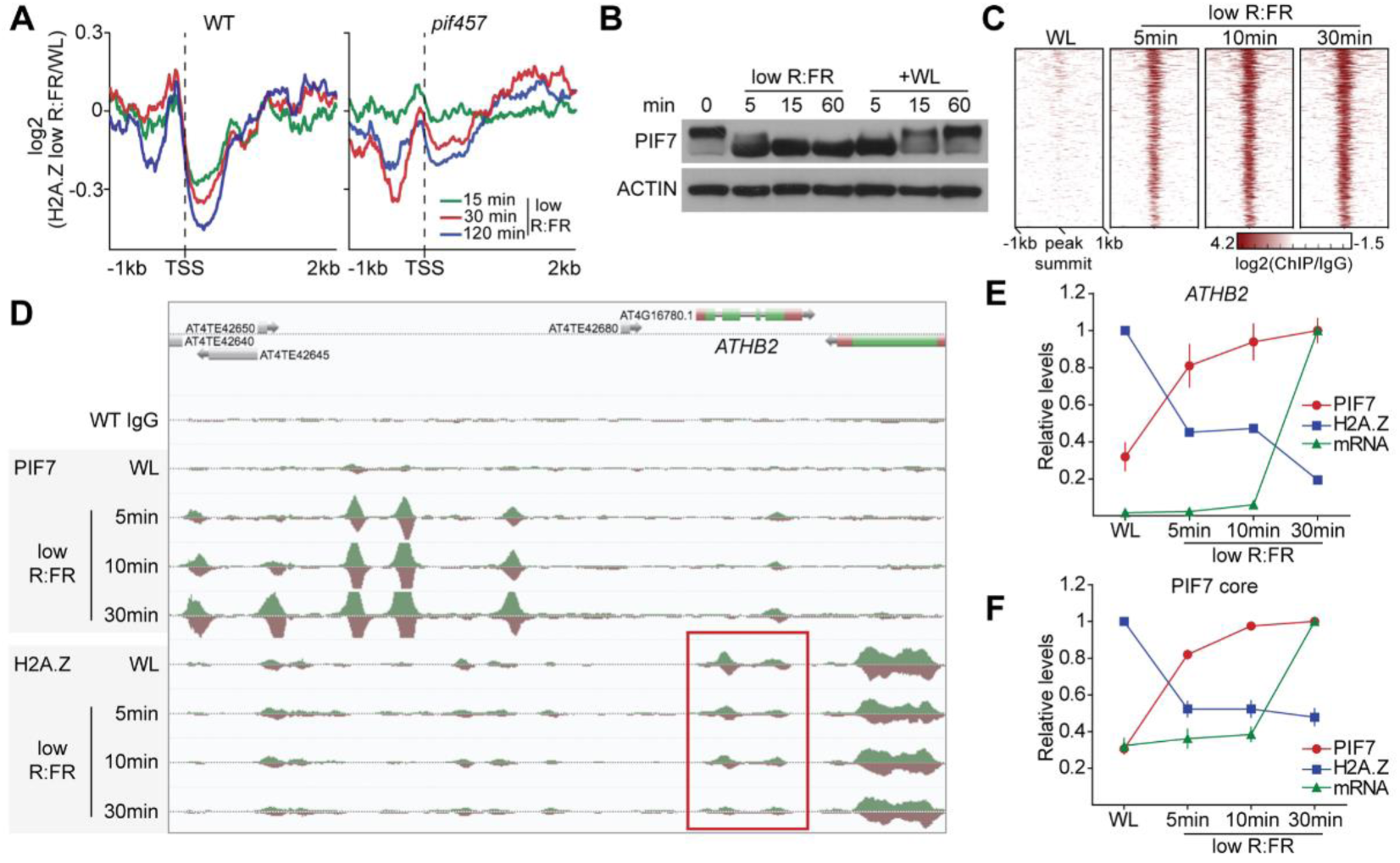
DNA binding of PIF7 initiates low R:FR-induced H2A.Z eviction at its target genes. (A) Aggregated profiles display low R:FR-induced H2A.Z eviction in WT and *pif457* seedlings. Profiles visualize the log_2_ fold change between WL and low R:FR-exposed H2A.Z ChIP-seq samples at the indicated lengths of low R:FR exposures. Eviction of H2A.Z is visualized from 1 kb upstream to 2 kb downstream of the transcriptional start site (TSS) of 200 genes that show strong H2A.Z eviction in WT seedlings after two hours of low R:FR light exposure. (B) Immunoblot analysis of *pif457 PIF7:PIF7:4xMYC* grown in constant WL. 6-day-old seedlings were exposed to low R:FR for 5, 15 and 60 min and subsequently moved back to WL for additional 5, 15 and 60 min. (C) Heatmap visualizes the low R:FR-induced DNA binding (log_2_ fold change) of PIF7 in *pif457 PIF7:PIF7:4xMYC* seedlings at the indicated time points. PIF7 binding is shown for the top 500 binding sites that display the strongest binding after 30 min of low R:FR exposure. PIF7 occupancy is shown from 1 kb upstream to 1 kb downstream of the 500 strongest PIF7 binding events. (D) AnnoJ genome browser screenshot visualizes PIF7 binding and H2A.Z occupancy at the *ATHB2* gene over time. Genome-wide occupancy of PIF7 and H2A.Z was determined in the same *pif457 PIF7:PIF7:4xMYC* chromatin by ChIP-seq. All tracks were normalized to the respective sequencing depth. The area marked in red indicates the gene body region of *ATHB2*. (E) (F) Quantification of relative PIF7 binding (red) and H2A.Z levels (blue) at *ATHB2* (E) and the PIF7 core gene set (F) is shown. PIF7 binding in peak summit regions was calculated as the ratio between two PIF7 ChIP-seq replicates sample and the WT MYC control. Levels of PIF7 binding at 30 min of low R:FR exposure were set to 1 for each target gene. H2A.Z occupancy for gene body-localized H2A.Z domains that were identified with SICER was calculated as the ratio between H2A.Z ChIP-seq sample and the WT IgG control. H2A.Z levels of WL-exposed *pif457 PIF7:PIF7:4xMYC* seedlings were set to 1. mRNA expression (green) is measured in TPM (Transcripts Per Kilobase Million) and is also derived from *pif457 PIF7:PIF7:4xMYC* seedlings. Expression values at 30 min of low R:FR exposure were set to 1 for each gene. The mean ± SEM represents PIF7 levels in the 5 binding peaks at *ATHB2* (E), and levels of PIF7 binding at the 20 PIF7 core genes (F). It also represents levels of mRNA expression of the 20 PIF7 core genes (F).

In humans, yeast and *Arabidopsis*, H2A.Z eviction is facilitated by the SWI/SNF-type ATP-dependent chromatin remodeler INO80 (*8, 20, 21*). In addition, the histone chaperone ANP32E was also reported to mediate H2A.Z eviction in humans (*22, 23*). While *anp32e* mutants displayed no phenotype, *ino80* mutants showed a significantly shorter hypocotyl after low R:FR exposure. Similar phenotypes were also observed in other INO80 complex subunit mutants including ARP5 (*arp5*) and EEN (*een*), as well as in *ino80 een* double mutants (Fig. 3A). We then tested diverse INO80 subunits (e.g. ARP5, EEN, RVB2) in pull-downs and discovered EEN as a PIF interacting protein (Fig. S5A) which was confirmed by further *in vitro* and *in vivo* studies (Fig. 3B, C). Moreover, the *een* mutant partially suppressed the PIF-dependent long hypocotyl phenotype of *phyB* (Fig. 3D) and overexpression of *EEN* was able to complement the *een* mutant hypocotyl phenotype (Fig. S5B). Since EEN, the *Arabidopsis* homologue of yeast Ies6, is part of the *Arabidopsis* INO80 complex which is involved in regulating H2A.Z eviction (*8*), we tested H2A.Z dynamics in *een* and *pif457* mutants at ZT4. Strikingly, the low R:FR-induced H2A.Z eviction was compromised in *een* and *pif457* mutants (Fig. 3E, Table S9), indicating that cooperative activity of PIFs and the INO80 complex directly shapes the global H2A.Z landscape in response to changes in light quality (Fig. 3F).

**Fig. 3.**
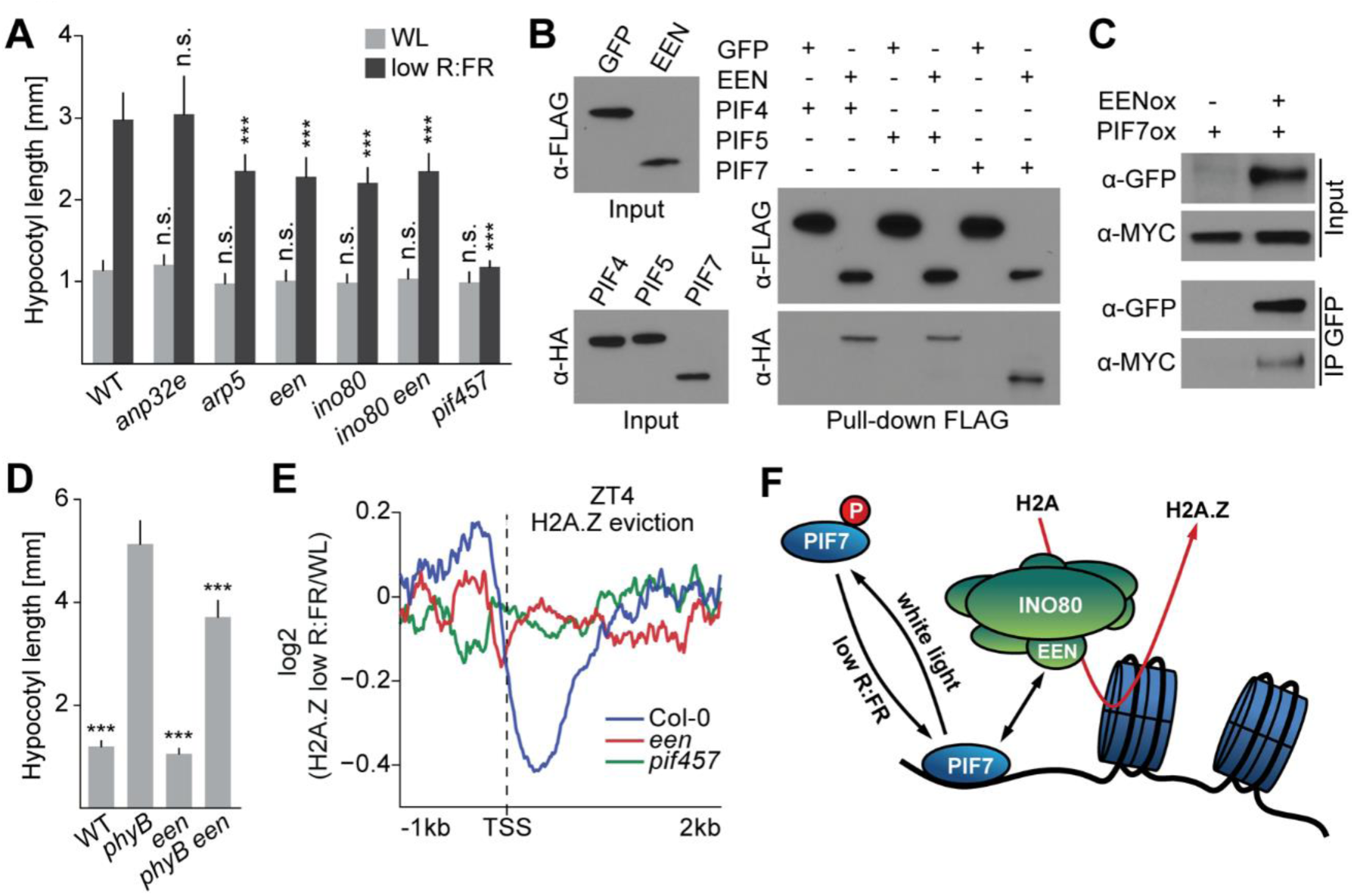
A PIF7-INO80 regulatory module facilitates low R:FR-induced H2A.Z eviction. (A) Hypocotyl length measurements of WT and mutant seedlings grown in WL or in responses to low R:FR. Error bars for the hypocotyl length represent SD (n ≥ 20). Stars denote statistically significant differences between WT and mutants grown in WL or WT and mutant grown in low R:FR, respectively (Two-way ANOVA, Tukey’s multiple comparisons test, n.s. P > 0.05 * P ≤ 0.05, ** P ≤ 0.01, *** P ≤ 0.001). (B) Hypocotyl length measurements of WT, *phyB*, *een* and *phyB een* grown in WL. Error bars for the hypocotyl length represent SD (n ≥ 32). Stars denote statistically significant differences between *phyB* and the other genotypes (One-way ANOVA, Tukey’s multiple comparisons test, n.s. P > 0.05 * P ≤ 0.05, ** P ≤ 0.01, *** P ≤ 0.001). (C) Pull-down assay using *in vitro* translated proteins. EEN was tagged with FLAG and PIFs with HA. FLAG:GFP served as a negative control. (D) Co-IP using 6-day-old seedlings overexpressing MYC-tagged PIF7 alone or in combination with GFP-tagged EEN. (E) Aggregated profiles display low R:FR-induced H2A.Z eviction in WT, *een* and *pif457* seedlings at ZT4. Profiles visualize the log_2_ fold change between WL and low R:FR-exposed H2A.Z ChIP-seq samples. Eviction of H2A.Z is visualized from 1 kb upstream to 2 kb downstream of the TSS of 464 genes that show H2A.Z eviction (≥ 1.25) fold after low R:FR exposure in WT seedlings. (F) Model for the control of low R:FR-induced H2A.Z eviction through the PIF7-INO80 regulatory module. Upon exposure to low R:FR light, PIF7 gets dephosphorylated and subsequently binds to its target sites where it potentially interacts with the EEN subunit of the INO80 complex. Consequently, gene body-localized H2A.Z gets evicted and gene expression of PIF7 target genes will be initiated.

## Discussion

By exploiting the rapid low R:FR light-mediated activation of PIF7 activity, we simultaneously captured global changes in PIF7 binding and H2A.Z occupancy at high temporal resolution. We provide strong evidence that although the low R:FR-induced DNA binding of PIF7 and H2A.Z eviction at PIF7 target genes showed similar kinetics, H2A.Z eviction requires PIF DNA binding. This is in contrast to the reported function of H2A.Z eviction at the *FT* locus in high ambient temperatures where it mediates PIF4 binding directly (*4*). Our finding of a PIF4-EEN interaction suggests that PIF4 can also remove H2A.Z through the INO80 complex. Thus, we speculate that the chromatin dynamics observed at the *FT* locus may represent a rather unique case. We utilized light quality changes to identify the PIF-INO80 regulatory module. Considering the functional diversity of PIFs, our findings are potentially translatable to other agronomically important responses and will pave the way for future investigations of plant-environment interactions which is critically needed in times of climate change.

## Acknowledgements

We thank Dr. Xuelin Wu for materials and advice regarding Gibson cloning and Tisha Haque for help with genotyping.

## Funding

B.C.W. was supported by an EMBO Long-Term Fellowship (ALTF 1514-2012), the Human Frontier Science Program (LT000222/2013-L) and the Salk Pioneer Postdoctoral Endowment Fund. M.Z. was supported by the Salk Pioneer Postdoctoral Endowment Fund as well as by a Deutsche Forschungsgemeinschaft (DFG) research fellowship (Za-730/1-1). This work was supported by grants from the National Science Foundation (NSF) (MCB-1024999, to J.R.E.), the Division of Chemical Sciences, Geosciences, and Biosciences, Office of Basic Energy Sciences of the U.S. Department of Energy (DE-FG02-04ER15517, to J.R.E.), the Gordon and Betty Moore Foundation (GBMF3034, to J.R.E.) and the National Institutes of Health (NIH) (5R35 GM122604, to J.C.). J.C. and J.R.E. are investigators of the Howard Hughes Medical Institute.

## Contributions

B.C.W., M.Z., J.R.E. and J.C. designed the research. B.C.W. and M.Z. performed RNA-seq and ChIP-seq experiments. M.Z., Y.H., J.R.N. and H.C. analyzed the sequencing data and performed bioinformatics analyses. Plasmid cloning was done by B.C.W., M.Z. and A.P. Generation of genetic material, phenotyping, western blotting, pull-down and Co-IP experiments were conducted by B.C.W and by A.P. under B.C.W.’s supervision. R.G. and S.T. shared plasmid clones and protein interaction data. B.C.W., M.Z., J.R.E. and J.C. prepared the figures and wrote the manuscript.

## Competing interests

Authors declare no competing interests.

---

**Fig. S1.**
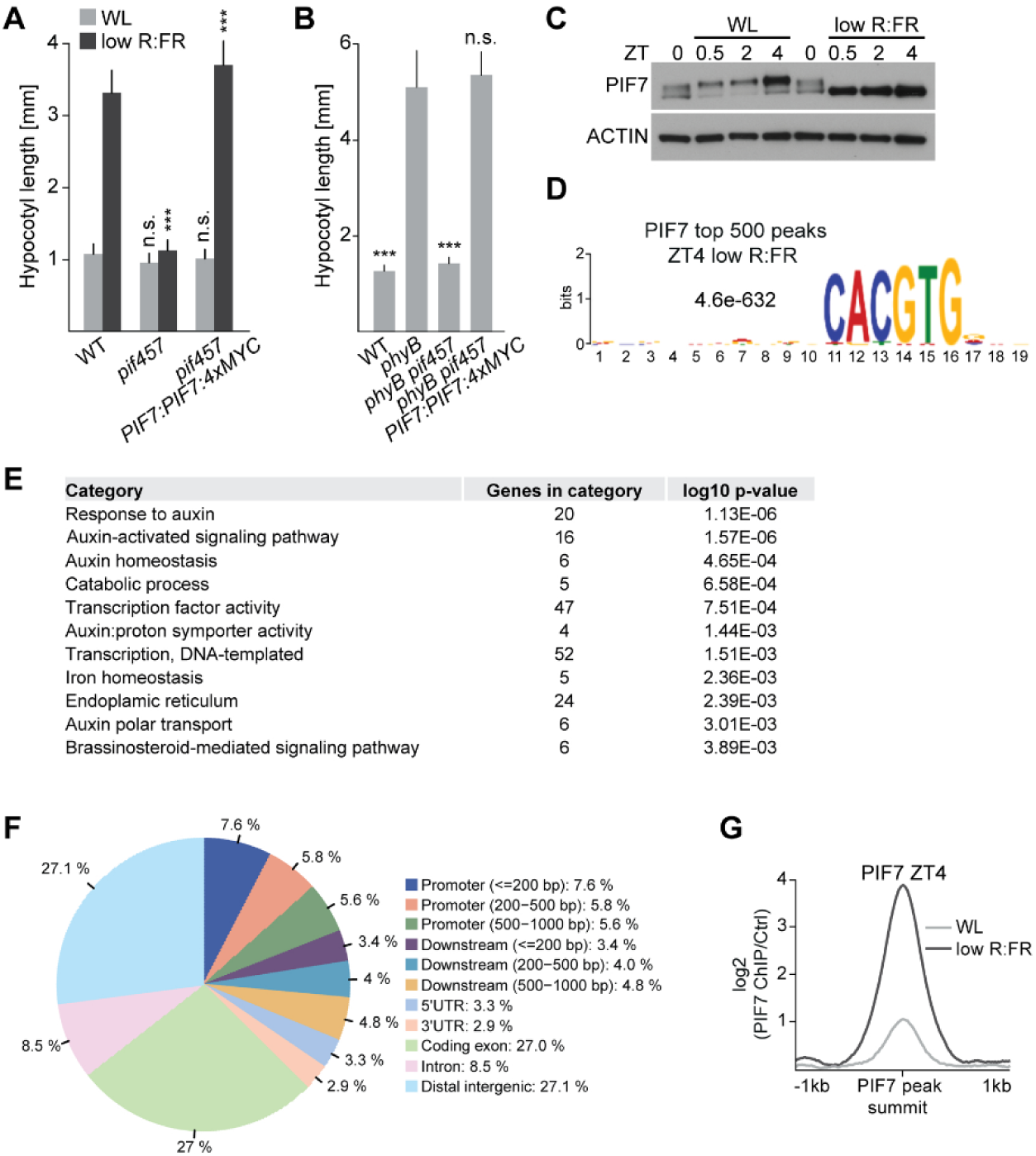
Low R:FR light exposure induces global PIF7 DNA binding. (A) Hypocotyl length measurements of WT, *pif457* and *pif457* expressing *PIF7:PIF7:4xMYC* in white light or in responses to low R:FR. Error bars for the hypocotyl length represent SD (n ≥ 30). Stars denote statistically significant differences between WT and the other genotypes grown in WL or WT and the other genotypes grown in low R:FR, respectively (Two-way ANOVA, Tukey’s multiple comparisons test, n.s. P > 0.05 * P ≤ 0.05, ** P ≤ 0.01, *** P ≤ 0.001). (B) Hypocotyl length measurements of WT, *phyB*, *phyB pif457* and *phyB pif457* expressing *PIF7:PIF7:4xMYC* grown in WL. Error bars for the hypocotyl length represent SD (n ≥ 14). Stars denote statistically significant differences between *phyB* and the other genotypes (One-way ANOVA, Tukey’s multiple comparisons test, n.s. P > 0.05 * P ≤ 0.05, ** P ≤ 0.01, *** P ≤ 0.001). (C) Immunoblot analysis of *pif457 PIF7:PIF7:4xMYC* in LD. 6-day-old seedlings continued to grow in WL or where exposed to low R:FR at ZT0. (D) The CACGTG motif was the top-ranked motif in PIF7 ChIP-seq derived from low R:FR-exposed *pif457 PIF7:PIF7:4xMYC* seedlings at ZT4. Motif discovery was done through MEME analysis using the 500 top-ranked peaks. (E) Gene ontology enrichment analysis revealed an enrichment for auxin signaling in the 500 top-ranked PIF7 targets. (F) The genomic distribution of PIF7 binding at ZT4 after low R:FR exposure is illustrated as pie chart. (G) Aggregated profile shows the low R:FR dependent difference between PIF7 binding at ZT4. PIF7 binding was determined in WL and low R:FR-exposed *pif457 PIF7:PIF7:4xMYC* seedlings by ChIP-seq. PIF7 occupancy is shown from 1 kb upstream to 1 kb downstream of the 500 strongest PIF7 binding events.

**Fig. S2.**
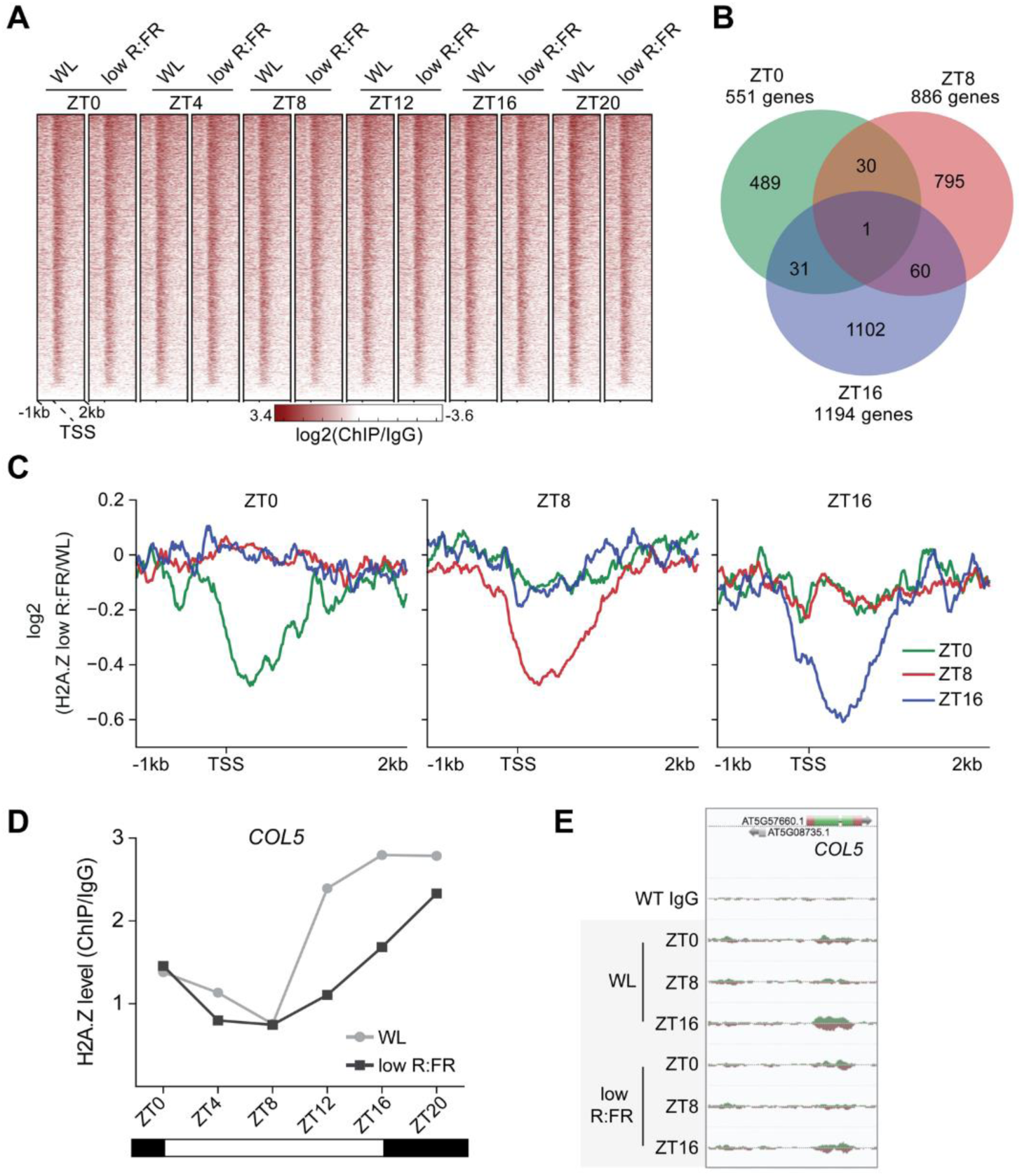
Low R:FR light exposure and the diurnal cycle control genome-wide H2A.Z occupancy in a daytime-dependent manner. (A) Heatmap visualizes absolute H2A.Z of all *Arabidopsis thaliana* protein-coding genes (TAIR10) at the indicated time points and light treatments. H2A.Z occupancy was determined by ChIP-seq in WT seedlings and calculated as the log_2_ fold change between H2A.Z ChIP and IgG control sample. (B) Venn diagram shows overlap between genes that show low R:FR-induced H2A.Z eviction (≥ 1.3 fold) at ZT0, ZT8 and ZT16. (C) Aggregated profiles visualize low R:FR-induced H2A.Z eviction at ZT0, ZT8 and ZT16 in WT seedlings. Eviction of H2A.Z is visualized from 1 kb upstream to 2 kb downstream of the TSS. (D) Quantification of H2A.Z levels at the gene body of *COL5* is shown. Occupancy of H2A.Z was determined by ChIP-seq and calculated as the ratio between H2A.Z and IgG control. (E) AnnoJ genome browser screenshot visualizes the light quality-dependent H2A.Z occupancy at the *COL5* gene at ZT0, ZT8 and ZT16. The WT IgG track serves as a control and all tracks were normalized to their sequencing depth.

**Fig. S3.**
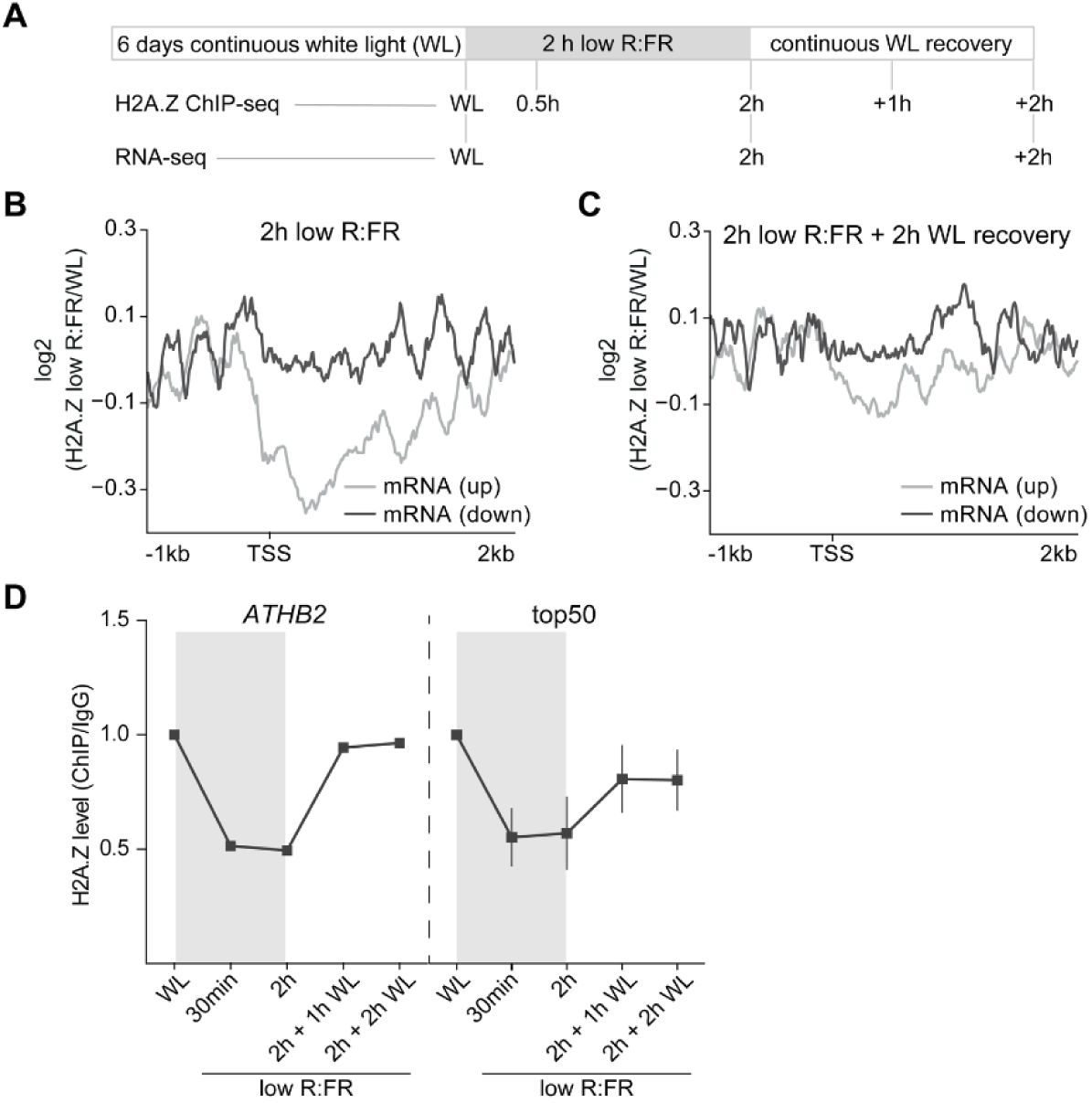
Low R:FR light-mediated manipulation of H2A.Z dynamics. (A) Schematic overview illustrates the experimental setup that was used to investigate chromatin dynamics in low R:FR light responses. (B) (C) Aggregated profiles visualize low R:FR-induced H2A.Z eviction and incorporation after two hours of low R:FR exposure (B) and after an additional two-hour-long WL recovery phase (C). Profiles are shown for genes that are differentially expressed after two hours of low R:FR exposure (blue for up-regulated genes and red for down-regulated genes). (D) Quantification of H2A.Z levels at the gene body of *ATHB2* and the 50 genes (top50) that show the strongest H2A.Z eviction after two hours of low R:FR exposure. Occupancy of H2A.Z was determined by ChIP-seq and calculated as the ratio between H2A.Z and IgG. H2A.Z levels of WL-exposed seedlings were set to 1.

**Fig. S4.**
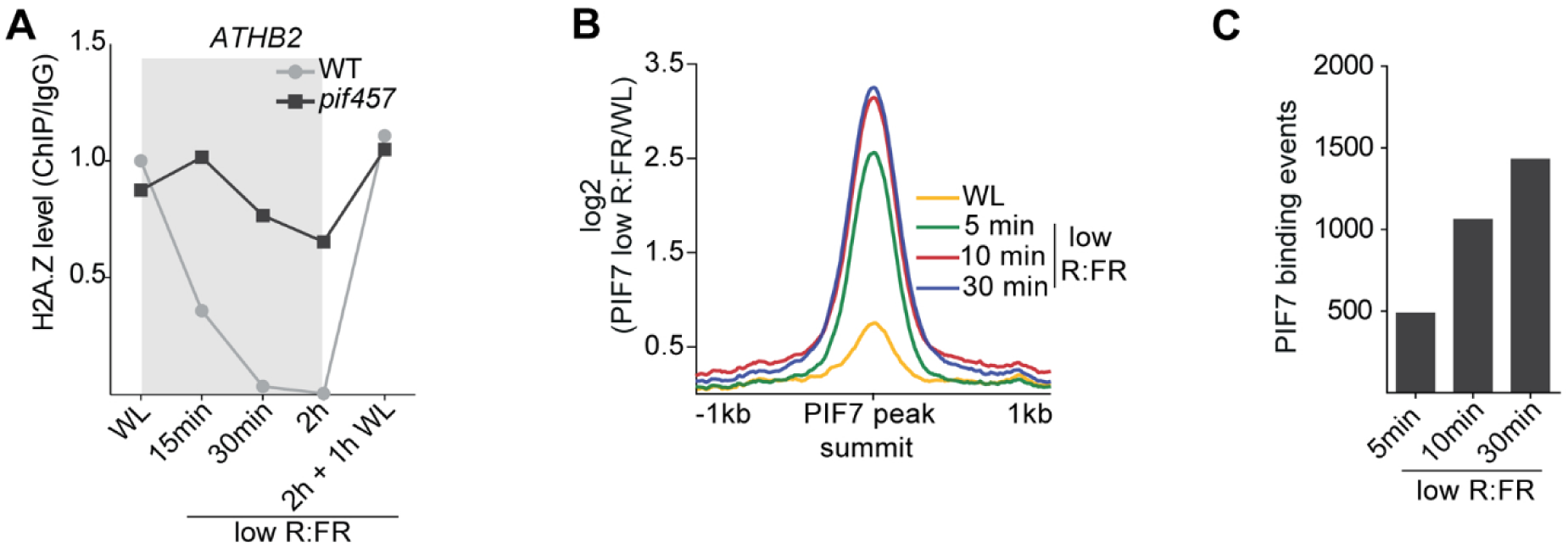
Low R:FR-induced PIF7 DNA binding is required for H2A.Z eviction. (A) Levels of H2A.Z at *ATHB2* in WT and *pif457* seedlings at the indicated time points are shown. Occupancy of H2A.Z was determined by ChIP-seq and calculated as the ratio between H2A.Z and IgG. (B) Aggregated profiles visualize the low R:FR-mediated activation of PIF7 after short low R:FR exposures (5, 10 and 30 min). PIF7 binding was determined in WL and low R:FR-exposed *pif457 PIF7:PIF7:4xMYC* seedlings by ChIP-seq and was calculated as the ratio between H2A.Z ChIP-seq samples and IgG control sample. PIF7 occupancy is shown from 1 kb upstream to 1 kb downstream of the 500 strongest PIF7 binding events. (C) Bar plot illustrates increase of low R:FR-induced PIF7 DNA binding events. PIF7 binding events were determined through the direct comparison of the respective low R:FR-exposed and WL-exposed PIF7 ChIP-seq replicates.

**Fig. S5.**
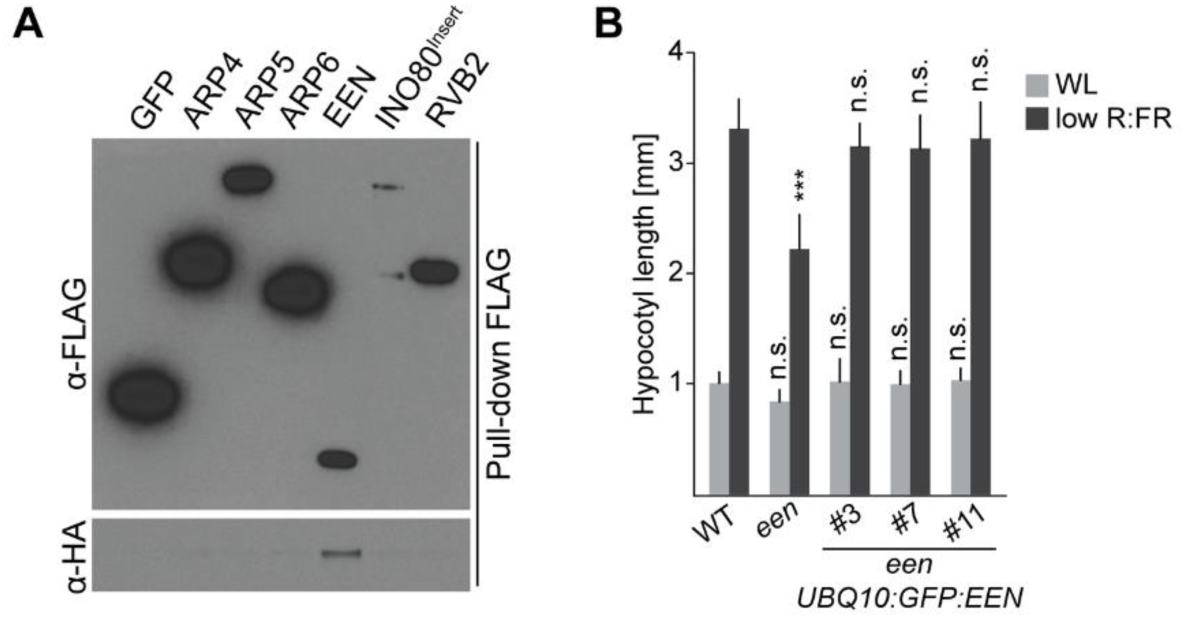
PIF-EEN interaction and *een* mutant complementation. (A) Pull-down assay with *in vitro* translated proteins. ARP4, ARP5, ARP6, EEN, INO80 insertion domain (INO80^Insert^) (*24*), and RVB2 were tagged with FLAG and PIF4 with HA. FLAG:GFP served as negative control. (B) Hypocotyl length measurements of WT, *een* and three *een UBQPIF7:PIF7:4xMYC* lines. Error bars for the hypocotyl length represent SD (n ≥ 15). Stars denote statistically significant differences in comparison to WT. WL was compared to WL and low R:FR to low R:FR (Two-way ANOVA, Tukey’s multiple comparisons test, n.s. P > 0.05 * P ≤ 0.05, ** P ≤ 0.01, *** P ≤ 0.001).

## Materials and Methods

### Genetic material and plasmid cloning

All genetic material used in this study is in the *Arabidopsis* Columbia-0 (Col-0) background (WT). *pif4-101 (25)*, *pif5-2* (*26*), *pif5-3* (*26*), *pif7-1* (*27*), *pif4-101 pif5-3* (*25*), *pif4-2 pif7-1* (*27*), *pif1-1 pif3-3 pif4-2 pif5-3* (*28*), *phyB-9* (*29*), *phyB-9 pif4-101* (*25*), *phyB-9 pif7-1* (*27*), *arp5-1* (*30*), *een-2* (*8*) and *ino80-8* (*8*) were described previously. By crossing these lines following mutant combinations were generated for this study: *pif5-3 pif7-1*, *pif4-2 pif5-3 pif7-1*, *pif1-1 pif3-3 pif4-2 pif5-3 pif7-1*, *phyB-9 pif5-2*, *phyB-9 pif4-101 pif5-2, phyB-9 pif4-101 pif7-1*, *phyB-9 pif5-2 pif7-1*, *phyB-9 pif4-101 pif5-2 pif7-1* and *ino80-8 een-2*. The T-DNA insertion in AT3G50690 (homologue of human ANP32E) is located in the first exon (SALK_033316).

For *PIF7:PIF7:4xMYC*, a 4064 bp genomic *PIF7* fragment was amplified by PCR and integrated via Gateway cloning and pDONR221 (Thermo Fisher Scientific) into pGWB16 (*31*). For *UBQ10:GFP:EEN*, the coding sequence from *EEN* in pDONR221 was integrated into pGWB6 (*31*) and *GFP:EEN* was amplified by PCR. Additionally, UBQ10 promoter and RCBS terminator were amplified from UBQ10pro in pDONR-P4P1R (*32*) or pMX202 (*33*), respectively. Promoter, coding sequence and terminator were cloned into the SmaI site of pJHA212G (*34*) via Gibson Assembly cloning (NEB). Floral dip was performed for all *Arabidopsis* transformations (*35*). *PIF7:PIF7:4xMYC* was transformed into *pif4-101 pif5-3 pif7-1* and homozygous T3 lines were crossed with *phyB-9 pif4-101 pif5-2 pif7-1*. *UBQ10:GFP:EEN* was transformed into *een-2* and into *pif7-2 35S:PIF7:9xMyc:6xHis:3xFlag* (*10*). For *in vitro* transcription and translation, coding sequences were derived from *Arabidopsis* cDNA or from the AtORFeome2.0 clone collection and were Gateway cloned into pTnT FLAG:GW or pTnT HA:GW (*36*). Primer sequences can be found in Table S10.

### Growth conditions

Seedlings were grown on half-strength Linsmaier and Skoog media in LED chambers (Percival Scientific) at 21ºC in long day conditions (16 h day/8 h night) or constant light. For white light conditions, white LED light was supplemented with red LED light, yielding a photosynthetically active radiation (PAR) of 110 μmol⋅m^−2^⋅s^−1^. For low R:FR conditions, far-red LEDs were used to decrease the R:FR ratio to 0.6, while fluence rates for all other wavelengths and therefore PAR were unchanged in comparison to white light conditions.

### Hypocotyl measurement

Seeds of experiments using *arp5*-1, *een*-2 and *ino80*-8 seeds were stratified for at least 5 days, otherwise, seeds were stratified for a minimum of 3 days. Afterwards, plates were exposed for 5 hours to 30 μmol⋅m^−2^⋅s^−1^ red light, transferred to darkness for 19 hours to synchronize germination and grown in LD. 9 days after stratification agar plates were scanned and hypocotyls were measured using NIH ImageJ software. For low R:FR treatment, seedlings were moved from white light to low R:FR light conditions 5 days after stratification.

### Western blotting

Frozen 6-day-old seedlings were disrupted in a ball mill using glass beads. Ground tissue was boiled in 2X NuPAGE LDS Sample Buffer (including 1.8% β-Mercaptoethanol) for 5min and separated in NuPAGE 4-12% Bis-Tris Protein Gels (Thermo Fisher Scientific). Following antibodies were used for immunoblotting: Myc-Tag, 9B11 (2276, Cell Signaling Technology), Anti-HA-Peroxidase, High Affinity clone 3F10 (11867423001, Roche), Anti-FLAG M2-Peroxidase (HRP) Clone M2 (A8592, MilliporeSigma), anti-GFP clones 7.1 & 13.1 (11814460001, Roche) and Goat Anti-Mouse IgG (H+L)-HRP Conjugate (1706516, Bio-Rad).

### Protein pull-down assay

According to manufacturer’s (Promega) instructions, TNT SP6 Coupled Wheat Germ Extract System was used to express HA fusions, while TNT SP6 Coupled Reticulocyte Lysate System served to express FLAG fusions. All further steps were performed at 4 °C. Reaction mixes were diluted with Paca buffer I (50 mM Tris pH 7.5, 100 mM NaCl, 1 mM EDTA, 1 mM TCEP pH 7.5, 1% (v/v) DMSO, 0.1% (v/v) IGEPAL CA-630, 0.04% (v/v) Tween-20) and FLAG-tagged protein were incubated for 1 hr with Anti-FLAG M2 Affinity Gel (Sigma) while rotating. Beads were washed three times with Paca buffer I and HA-tagged proteins were added. After rotating for 30 min, beads were washed four times with Paca buffer I, boiled in 2X NuPAGE LDS Sample Buffer (including 1.8% β-Mercaptoethanol) for 5min and separated in NuPAGE 4-12% Bis-Tris Protein Gels (Thermo Fisher Scientific).

### Protein Co-Immunoprecipitation

At ZT0, 6-day-old seedlings were exposed to low R:FR for 8 hours. Frozen material was disrupted in a ball mill using glass beads. Plant material was resuspended in Paca buffer II+ (50 mM Tris pH 7.5, 100 mM NaCl, 1 mM EDTA, 1 mM TCEP pH 7.5, 1% (v/v) DMSO, 0.4% (v/v) IGEPAL CA-630, 1X cOmplete Protease Inhibitor Cocktail, Roche, 125 U/ml Benzonase, MilliporeSigma) and rotated for 30 min at 4 °C. Supernatant was cleared by centrifugation and incubated for 1 hr with ChromoTek GFP-Trap. Beads were washed six times with Paca buffer II-(excluding Benzonase), boiled in 2X NuPAGE LDS Sample Buffer (including 1.8% β-Mercaptoethanol) for 5 min and separated in NuPAGE 4-12% Bis-Tris Protein Gels (Thermo Fisher Scientific).

### ChIP-sequencing

With the exception of the experiment depicted in Fig. S2, seedlings were crosslinked 6 days after stratification. For the experiment shown in Fig. S2, 5 days after stratification, seedlings continued to grow in WL or were moved to low R:FR conditions. Samples were crosslinked 8 days after stratification. ChIP-seq experiments were performed as previously described (*37*) with minor modifications. ChIP-seq assays were conducted with antibodies against H2A.Z (39647, Active Motif) and MYC-tag Mouse (2276, Cell Signaling Technology). IgG (015-000-003, Jackson ImmunoResearch) served as the negative control. Dynabeads Protein G (Thermo Fisher Scientific) were coupled for 4–6 hours with respective antibodies and incubated overnight with equal amounts of sonicated chromatin. Beads were consecutively washed with high salt buffer (50 mM Tris HCl pH 7.4, 150 mM NaCl, 2 mM EDTA, 0.5% Triton X-100), low salt (50 mM Tris HCl pH 7.4, 500 mM NaCl, 2 mM EDTA, 0.5% Triton X-100) and wash buffer (50 mM Tris HCl pH 7.4, 50 mM NaCl, 2 mM EDTA) before carrying out de-crosslinking at 65 ºC, Proteinase K treatment and DNA precipitation. Libraries were sequenced on Illumina HiSeq 2500 and HiSeq 4000 Sequencing systems. Sequencing reads were aligned to TAIR10 genome assembly using Bowtie2 (*38*).

### RNA-sequencing

Total RNA was extracted using RNeasy Plant Mini Kit (Qiagen). cDNA library preparation and single read sequencing was performed as described previously (*39*). Sequencing reads were aligned to TAIR10 genome assembly using STAR software (STAR_2.6.0 c) (*40*).

### Sequencing data analysis

H2A.Z occupancy was determined from ChIP-seq experiments with the SICER software (*41*) using the TAIR10 genome assembly and WT IgG samples as a control. Genes that were most proximal to H2A.Z enriched domains were identified with the Intersect tool from BEDtools (*42*). For the identification of PIF7 peak summit regions, we used the Genome wide Event finding and Motif discovery (GEM) tool (version 2.5) (*43*). WT chromatin treated with an anti-MYC antibody served as control for the total number of PIF7 ChIP-seq peaks. For the low-R:FR-specific identification of PIF7 peaks, we used chromatin from WL-exposed *pif457 PIF7:PIF7:4xMYC* seedlings as a control. We used the MEME-ChIP analysis tool (*44*) to identify preferred binding motifs within the top 500 summit regions of two merged PIF7 ChIP-seq experiments at ZT4 with low R:FR exposure. Biological ChIP-seq replicates were merged with SAMtools (*45*). DAVID was used to identify gene ontology (GO) enrichment in the PIF7 ChIP-seq data (*46*). For the analysis of genomic distributions within PIF7 ChIP-seq data the *cis*-regulatory element annotation system (CEAS) tool was used (*47*). Heatmaps and aggregated profiles of ChIP-seq data were carried out with Deeptools (*48*). To quantify occupancy of H2A.Z and PIF7 at *ATHB2* and the PIF7 core gene set, we employed the bigWigAverageOverBed tool executable from the UCSC genome browser (*49*). For the identification of genes with a low R:FR-induced H2A.Z eviction, we employed SICER. Transcripts in the RNA-seq data were quantified with the RSEM software package (version 1.3.0) and differentially induced genes were identified with the Cufflinks package (*50*). The AnnoJ genome browser was used to visualize all sequencing data (*51*).

## Supplementary Tables

**Table S1** List of significant PIF7 DNA binding events at ZT4 under low R:FR light exposure

**Table S2** List of genes with significant H2A.Z enrichment throughout a LD time course

**Table S3** List of genes that show low R:FR-induced H2A.Z eviction at ZT0, ZT8 and ZT16

**Table S4** List of differentially expressed genes after 2 hours of low R:FR light exposure

**Table S5** List of genes that show H2A.Z eviction after 2 hours of low R:FR light exposure

**Table S6** List of top 200 genes with impaired R:FR light-induced H2A.Z eviction in *pif457* mutants

**Table S7** List of significant PIF7 DNA binding events after short low R:FR light exposure times

**Table S8** The PIF7 core gene set

**Table S9** List of genes that show low R:FR-induced H2A.Z eviction in WT seedlings at ZT4

**Table S10** List of primers used in this study

## References

1. K. A. Franklin, P. H. Quail, Phytochrome functions in Arabidopsis development. J Exp Bot 61, 11–24 (2010).

2. K. A. Franklin, G. Toledo-Ortiz, D. E. Pyott, K. J. Halliday, Interaction of light and temperature signalling. J Exp Bot 65, 2859–2871 (2014).

3. P. Leivar, E. Monte, PIFs: systems integrators in plant development. Plant Cell 26, 56–78 (2014).

4. S. V. Kumar et al., Transcription factor PIF4 controls the thermosensory activation of flowering. Nature 484, 242–245 (2012).

5. S. V. Kumar, P. A. Wigge, H2A.Z-containing nucleosomes mediate the thermosensory response in Arabidopsis. Cell 140, 136–147 (2010).

6. W. Sura et al., Dual Role of the Histone Variant H2A.Z in Transcriptional Regulation of Stress-Response Genes. Plant Cell 29, 791–807 (2017).

7. D. Coleman-Derr, D. Zilberman, Deposition of histone variant H2A.Z within gene bodies regulates responsive genes. PLoS Genet 8, e1002988 (2012).

8. M. Zander et al., Epigenetic silencing of a multifunctional plant stress regulator. Elife 8, (2019).

9. J. J. Casal, Shade avoidance. Arabidopsis Book 10, e0157 (2012).

10. L. Li et al., Linking photoreceptor excitation to changes in plant architecture. Genes Dev 26, 785–790 (2012).

11. Y. Zhang et al., A quartet of PIF bHLH factors provides a transcriptionally centered signaling hub that regulates seedling morphogenesis through differential expression-patterning of shared target genes in Arabidopsis. PLoS Genet 9, e1003244 (2013).

12. P. Hornitschek et al., Phytochrome interacting factors 4 and 5 control seedling growth in changing light conditions by directly controlling auxin signaling. Plant J 71, 699–711 (2012).

13. E. Oh et al., Genome-wide analysis of genes targeted by PHYTOCHROME INTERACTING FACTOR 3-LIKE5 during seed germination in Arabidopsis. Plant Cell 21, 403–419 (2009).

14. E. Oh, J. Y. Zhu, Z. Y. Wang, Interaction between BZR1 and PIF4 integrates brassinosteroid and environmental responses. Nat Cell Biol 14, 802–809 (2012).

15. C. Steindler et al., Shade avoidance responses are mediated by the ATHB-2 HD-zip protein, a negative regulator of gene expression. Development 126, 4235–4245 (1999).

16. G. Sessa et al., A dynamic balance between gene activation and repression regulates the shade avoidance response in Arabidopsis. Genes Dev 19, 2811–2815 (2005).

17. D. Coleman-Derr, D. Zilberman, Deposition of histone variant H2A.Z within gene bodies regulates responsive genes. PLoS Genet 8, e1002988 (2012).

18. W. Sura et al., Dual Role of the Histone Variant H2A.Z in Transcriptional Regulation of Stress-Response Genes. Plant Cell 29, 791–807 (2017).

19. M. Hassidim, Y. Harir, E. Yakir, I. Kron, R. M. Green, Over-expression of CONSTANS-LIKE 5 can induce flowering in short-day grown Arabidopsis. Planta 230, 481–491 (2009).

20. M. Papamichos-Chronakis, S. Watanabe, O. J. Rando, C. L. Peterson, Global regulation of H2A.Z localization by the INO80 chromatin-remodeling enzyme is essential for genome integrity. Cell 144, 200–213 (2011).

21. H. E. Alatwi, J. A. Downs, Removal of H2A.Z by INO80 promotes homologous recombination. EMBO Rep 16, 986–994 (2015).

22. A. Obri et al., ANP32E is a histone chaperone that removes H2A.Z from chromatin. Nature 505, 648–653 (2014).

23. Z. Mao et al., Anp32e, a higher eukaryotic histone chaperone directs preferential recognition for H2A.Z. Cell Res 24, 389–399 (2014).

24. S. Eustermann et al., Structural basis for ATP-dependent chromatin remodelling by the INO80 complex. Nature 556, 386–390 (2018).

25. S. Lorrain, T. Allen, P. D. Duek, G. C. Whitelam, C. Fankhauser, Phytochrome-mediated inhibition of shade avoidance involves degradation of growth-promoting bHLH transcription factors. Plant J 53, 312–323 (2008).

26. R. Khanna et al., The basic helix-loop-helix transcription factor PIF5 acts on ethylene biosynthesis and phytochrome signaling by distinct mechanisms. Plant Cell 19, 3915–3929 (2007).

27. P. Leivar et al., The Arabidopsis phytochrome-interacting factor PIF7, together with PIF3 and PIF4, regulates responses to prolonged red light by modulating phyB levels. Plant Cell 20, 337–352 (2008).

28. P. Leivar et al., Multiple phytochrome-interacting bHLH transcription factors repress premature seedling photomorphogenesis in darkness. Curr Biol 18, 1815–1823 (2008).

29. J. W. Reed, P. Nagpal, D. S. Poole, M. Furuya, J. Chory, Mutations in the gene for the red/far-red light receptor phytochrome B alter cell elongation and physiological responses throughout Arabidopsis development. Plant Cell 5, 147–157 (1993).

30. M. K. Kandasamy, E. C. McKinney, R. B. Deal, A. P. Smith, R. B. Meagher, Arabidopsis actin-related protein ARP5 in multicellular development and DNA repair. Dev Biol 335, 22–32 (2009).

31. T. Nakagawa et al., Development of series of gateway binary vectors, pGWBs, for realizing efficient construction of fusion genes for plant transformation. J Biosci Bioeng 104, 34–41 (2007).

32. Y. Jaillais et al., Tyrosine phosphorylation controls brassinosteroid receptor activation by triggering membrane release of its kinase inhibitor. Genes Dev 25, 232–237 (2011).

33. X. Wu et al., Modes of intercellular transcription factor movement in the Arabidopsis apex. Development 130, 3735–3745 (2003).

34. S. Y. Yoo et al., The 35S promoter used in a selectable marker gene of a plant transformation vector affects the expression of the transgene. Planta 221, 523–530 (2005).

35. S. J. Clough, A. F. Bent, Floral dip: a simplified method for Agrobacterium-mediated transformation of Arabidopsis thaliana. Plant J 16, 735–743 (1998).

36. K. Nito, C. C. Wong, J. R. Yates, 3rd, J. Chory, Tyrosine phosphorylation regulates the activity of phytochrome photoreceptors. Cell Rep 3, 1970–1979 (2013).

37. K. Kaufmann et al., Chromatin immunoprecipitation (ChIP) of plant transcription factors followed by sequencing (ChIP-SEQ) or hybridization to whole genome arrays (ChIP-CHIP). Nat Protoc 5, 457–472 (2010).

38. B. Langmead, Aligning short sequencing reads with Bowtie. Curr Protoc Bioinformatics Chapter 11, Unit 11 17 (2010).

39. L. Song et al., A transcription factor hierarchy defines an environmental stress response network. Science 354, (2016).

40. A. Dobin et al., STAR: ultrafast universal RNA-seq aligner. Bioinformatics 29, 15–21 (2013).

41. C. Zang et al., A clustering approach for identification of enriched domains from histone modification ChIP-Seq data. Bioinformatics 25, 1952–1958 (2009).

42. A. R. Quinlan, I. M. Hall, BEDTools: a flexible suite of utilities for comparing genomic features. Bioinformatics 26, 841–842 (2010).

43. Y. Guo, S. Mahony, D. K. Gifford, High resolution genome wide binding event finding and motif discovery reveals transcription factor spatial binding constraints. PLoS Comput Biol 8, e1002638 (2012).

44. P. Machanick, T. L. Bailey, MEME-ChIP: motif analysis of large DNA datasets. Bioinformatics 27, 1696–1697 (2011).

45. H. Li et al., The Sequence Alignment/Map format and SAMtools. Bioinformatics 25, 2078–2079 (2009).

46. G. Dennis, Jr. et al., DAVID: Database for Annotation, Visualization, and Integrated Discovery. Genome Biol 4, P3 (2003).

47. H. Shin, T. Liu, A. K. Manrai, X. S. Liu, CEAS: cis-regulatory element annotation system. Bioinformatics 25, 2605–2606 (2009).

48. F. Ramirez, F. Dundar, S. Diehl, B. A. Gruning, T. Manke, deepTools: a flexible platform for exploring deep-sequencing data. Nucleic Acids Res 42, W187–191 (2014).

49. W. J. Kent et al., The human genome browser at UCSC. Genome Res 12, 996–1006 (2002).

50. C. Trapnell et al., Transcript assembly and quantification by RNA-Seq reveals unannotated transcripts and isoform switching during cell differentiation. Nat Biotechnol 28, 511–515 (2010).

51. R. Lister et al., Highly integrated single-base resolution maps of the epigenome in Arabidopsis. Cell 133, 523–536 (2008).

